# Fully Hyperbolic Neural Networks: A Novel Approach to Studying Aging Trajectories

**DOI:** 10.1101/2024.10.01.616153

**Authors:** Hugo Ramirez, Davide Tabarelli, Arianna Brancaccio, Paolo Belardinelli, Elisabeth B. Marsh, Michael Funke, John C. Mosher, Fernando Maestu, Mengjia Xu, Dimitrios Pantazis

## Abstract

Characterizing age-related alterations in brain networks is crucial for understanding aging trajectories and identifying deviations indicative of neurodegenerative disorders, such as Alzheimer’s disease. In this study, we developed a Fully Hyperbolic Neural Network (FHNN) to embed functional brain connectivity graphs derived from magnetoencephalography (MEG) data into low dimensions on a Lorentz model of hyperbolic space. Using this model, we computed hyperbolic embeddings of the MEG brain networks of 587 individuals from the Cambridge Centre for Ageing and Neuroscience (Cam-CAN) dataset. Notably, we leveraged a unique metric—the radius of the node embeddings—which effectively captures the hierarchical organization of the brain, to characterize subtle hierarchical organizational changes in various brain subnetworks attributed to the aging process. Our findings revealed that a considerable number of subnetworks exhibited a reduction in hierarchy during aging, with some showing gradual changes and others undergoing rapid transformations in the elderly. Moreover, we demonstrated that hyperbolic features outperform traditional graph-theoretic measures in capturing age-related information in brain networks. Overall, our study represents the first evaluation of hyperbolic embeddings in MEG brain networks for studying aging trajectories, shedding light on critical regions undergoing significant age-related alterations in the large cohort of the Cam-CAN dataset.

## I. Introduction

A LZHEIMER’S disease (AD) is a global public health priority, particularly affecting older populations. Characterized by a gradual and relentless neurodegenerative process, AD includes a prolonged asymptomatic preclinical phase before the onset of mild cognitive impairment and significant cognitive decline. Remarkably, the complex brain processes, including amyloid deposition, tau phosphorylation, and neuroimaging changes, that lead to AD can begin more than two decades before clinical symptoms appear [1]. These early changes provide valuable opportunities to monitor shifts in brain health that may precede the onset of symptoms or the disease itself. Observations of these changes can be combined with established risk factors, such as age.

Recent breakthroughs in AD treatments have shown the greatest benefits in early-stage patients [2], making it crucial to assess early deviations from typical *aging trajectories* [3]. Early intervention could slow AD progression, offering hope for improved outcomes for those eventually affected by the disease. An excellent tool for establishing normal brain aging trajectories and identifying deviations is magnetoencephalography (MEG) [4], [5]. MEG offers exceptional temporal resolution, allowing for the characterization of subtle brain changes, especially in the frequency domain [6]. Additionally, MEG captures the fields produced by intraneuronal currents, providing a more direct index of neuronal activity than biomarkers measuring hemodynamic responses, such as fMRI or FDG-PET. With MEG, researchers can generate functional connectivity maps and study distinctive patterns of alterations in functional connectivity within large-scale brain systems [4].

Despite the complex relational structure of brain connectivity networks (or graphs), most existing MEG-based neuroimaging studies rely heavily on statistical analyses using traditional graph-theoretic measures, which are based on domain knowledge but may not be optimal for capturing complex hierarchical patterns [7], [8]. Commonly used measures include node degree, closeness centrality, betweenness centrality, global efficiency, and other measures assessed across preselected brain subnetworks, such as the default mode network and salience networks [9]. However, there is a pressing need for data-driven methods that can more effectively capture complex hierarchical structures while adapting to the data and *automatically* detecting features suitable for downstream tasks, such as modeling and characterizing brain aging trajectories. Recent advancements in neural network models have highlighted their significant potential in embedding complex human brain networks into latent spaces, where nodes are represented as low-dimensional vectors [10], [4], [5]. Typically, graph representation learning employs graph convolutional neural networks to map graph nodes into points within Euclidean space. However, Euclidean geometry often falls short due to its limited representational capacity, leading to high distortion when applied to brain networks. This distortion arises because brain networks are scale-free graphs with a hierarchical, tree-like structure, where the number of nodes increases exponentially as one moves from higher to lower hierarchical levels. This exponential growth exceeds the polynomial expansion capacity of Euclidean space, resulting in distorted embeddings [11].

In contrast, hyperbolic space, characterized by its negative curvature, effectively addresses the exponential growth of nodes in scale-free brain graphs [5]. The hyperbolic geometry expands exponentially as one moves away from the center, mirroring the growth patterns of brain networks. This approach offers several advantages for embedding brain networks, such as minimal distortion and preservation of both local and global geometric information. It enables faithful embeddings even in low-dimensional spaces, which benefits downstream tasks including the characterization of age trajectories, and the development of neural network models with lower complexity, higher generalization capacity, and reduced training data requirements [12], [13], [14].

A crucial property of hyperbolic embedding is the hierarchical organization of node embeddings: higher hierarchical regions are mapped near the center, while lower hierarchical regions are mapped towards the periphery [5]. Here, we leveraged this characteristic to delineate changes in the hierarchical structure of the brain across different ages. We designed a novel hyperbolic MEG brain network embedding framework that transforms high-dimensional, complex MEG brain networks into lower-dimensional hyperbolic representations. This involved creating and validating a new hyperbolic model based on the architecture of the fully hyperbolic neural network (FHNN) [15]. Using this model, we computed hyperbolic embeddings of the MEG brain networks of 587 individuals from the Cam-CAN dataset [16]. A unique metric—the radius of the node embeddings—was employed to effectively proxy the hierarchical organization of the brain. We utilized this metric to characterize subtle changes in the hierarchical organization of various brain subnetworks attributed to the aging process.

Our findings revealed that a considerable number of subnetworks exhibit a reduction in hierarchy during aging, with some displaying slow changes and others undergoing faster changes across age. Overall, our study presents the first evaluation of hyperbolic embeddings in MEG brain networks to study aging trajectories, revealing critical regions that undergo significant age-related alterations in the large cohort of the Cam-CAN dataset.

## II. Methods

### A. Participants, data acquisition, and pre-processing

The study cohort consisted of 587 healthy participants (age range 18–89 years; 295 Female; see Table I) from the Cam-CAN dataset [16]. All subjects were tested for the absence of serious neurological and psychiatric conditions as well as the absence of cognitive decline, determined by having a MiniMental State Examination (MMSE) score higher than 24 [17]. The subjects were drawn from the imaging component of the Cam-CAN study, which originally included approximately 700 individuals, each undergoing different imaging protocols. For our analysis, we selected participants with complete imaging data: MEG resting-state recordings, MEG emptyroom recordings (for noise covariance), and T1-weighted scans (for cortical anatomy). This initially yielded a dataset of 591 individuals, but 4 participants were excluded due to MRI segmentation errors, resulting in a final cohort of 587.

**TABLE 1.** Age Distribution of Cam-CAN Subjects.

| Group | Age Range | Number of Subjects | Female subjects |
| --- | --- | --- | --- |
| 1st Decade | 18 - 30 | 67 | 38 |
| 2nd Decade | 30 - 40 | 87 | 38 |
| 3rd Decade | 40 - 50 | 101 | 53 |
| 4th Decade | 50 - 60 | 85 | 42 |
| 5th Decade | 60 - 70 | 100 | 47 |
| 6th Decade | 70 - 80 | 92 | 53 |
| 7th Decade | 80 - 90 | 55 | 24 |
| Total Subjects | 18 - 90 | 587 | 295 |

Three-minute eyes-closed resting-state MEG recordings were collected for all participants using a 306-channel VectorView MEG system (102 magnetometers, 204 first-order planar gradiometers; sampling rate = 1000 Hz; high-pass filter = 0.03 Hz; low-pass filter = 330 Hz). Pre-processing was performed using the Fieldtrip software [18] and custom MATLAB code, focusing only on the gradiometer recordings (magnetometers were excluded due to overlapping information).

The MEG data were visually inspected and all channels exhibiting high levels of noise or SQUID jumps were excluded from analysis. The Signal Source Separation method (SSS [19]) was employed to suppress external magnetic interference, reconstruct bad channels, and compensate for head movements, as detected by the continuous head position monitoring system. To identify and discard segments contaminated by muscular activity, MEG channels filtered between 100 Hz and 140 Hz were transformed into z-scores relative to the entire channel time series. Segments with z-scores exceeding 5 for at least 200 milliseconds were removed from subsequent analysis. Subsequently, Independent Component Analysis (Infomax ICA [20]), was applied to data filtered between 0.5 Hz and 125 Hz. Components predominantly associated with cardiac (ECG) and electro-ocular (HEOG and VEOG) signals were identified and discarded.

### B. Source reconstruction

For each participant, anatomical MRI T1-weighted images were processed to extract models of the central surface, defined as the mid-line between the gray/white matter and the pial/gray interfaces. We used a standard CAT12^1^ anatomical pipeline to conduct bias normalization, denoising and skull-stripping, resulting in the central surface, the brain enclosing surface, and a high resolution surface modeling the participant head.

We then co-registered the MEG resting-state data to the subject-specific central surfaces by aligning both the anatomical landmarks and the isotrak points to the high resolution head surface. Subsequently, the central surface was decimated to the equivalent of a 5th order icosahedron, resulting in approximately 20,000 vertices with an average spatial resolution of 3.1 mm. This decimated surface served as the foundation for defining the source model for MEG source reconstruction, with dipole orientations modeled as fixed and perpendicular to the cortical mantle.

Utilizing the source model and the brain enclosing surface, a forward model was computed using the “single shell” method [21]. Subsequently, a Minimum Norm Estimate (MNE) inversion operator was computed using a pre-computed emptyroom noise covariance matrix, regularized at a 10% level [22]. Prior to this analysis, the empty-room data, available in the Cam-CAN dataset for each experimental session, underwent identical preprocessing procedures as the human resting-state data. For further insights into preprocessing and MNE inversion operator computation, refer to [23].

### C. MEG Functional Connectivity Estimation

For each participant, clean MEG resting state data were segmented in epochs of 2 seconds and projected to the central surface, using the pre-computed MNE inversion operator. Subsequently, the MEG sources were anatomically averaged onto 180 regions of interest (ROI) per hemisphere, totalling 360 ROIs of the Human Connectome Project Multi-Modal Parcellation atlas (HCP-MMP1)[24]. The averaging procedure involved identifying the predominant orientations of the dipoles within each ROI and aligning opposing dipoles through a process known as sign flipping to prevent spurious cancelations. This procedure yielded a single 2-second time series for each epoch per ROI, across all participants.

We computed the functional connectivity between ROIs in the alpha frequency band (8 - 13 Hz) using the Phase Locking Value (PLV) metric [25]:

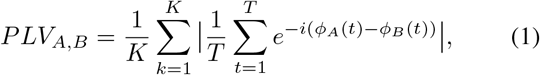

where *ϕ*_*A*_(*t*) and *ϕ*_*B*_(*t*) are the instantaneous phases of the signals *A* and *B* at the instant *t, T* is the total number of time points per epoch, and *K* is the total number of epochs. This computation yielded a 360 × 360 connectivity matrix for each of the 587 participants. We chose the alpha band because numerous studies have consistently reported abnormalities in alpha band oscillations in individuals with AD [26].

Finally, to convert the connectivity matrices into graphs appropriate for FHNN processing, we set a threshold of 0.360 on the PLV values, thereby keeping only the top 5% of edges and discarding those below this cutoff (Fig. 1). This threshold was chosen as a balanced trade-off to maintain the most significant connections in the brain networks while minimizing the impact of weaker ones. It’s important to note that including too many edges would result in overly dense brain networks, distorting the intrinsic scale-free properties of the original networks and reducing the benefits of hyperbolic embeddings.

**Fig. 1.**
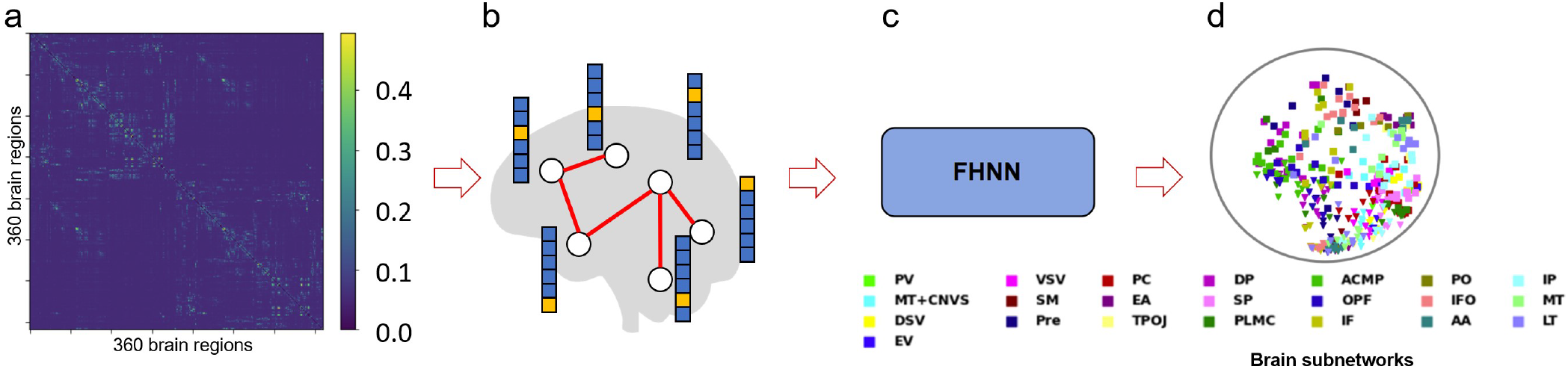
Hyperbolic embedding framework of MEG brain networks. (a) MEG functional connectivity matrices, calculated based on phase locking values across all 360 regions of the HCP-MMP1 brain atlas, were thresholded to keep 5% of the connectivity values. (b) Brain networks with the remaining connections are illustratively visualized together with corresponding one-hot node features. (c) The brain data were fed to a fully hyperbolic neural network, yielding embeddings for each brain region of interest. (d) Embedded nodes are color-coded based on membership in each of the 22 subnetworks with similar functional specialization as in [24]. Shape indicates hemisphere: “□” for right hemisphere, “**△** “ for left hemisphere. (PV: primary visual, VSV: ventral stream visual, PC: posterior cingulate, DP: dorsolateral prefrontal, ACMP: anterior cingulate and medial prefrontal, PO: posterior opercular IP: inferior parietal, MT+CNVS: MT + complex and neighboring visual areas, SM: somatosensory and motor, EA: early auditory SP: superior parietal, OPF: orbital and polar frontal, IFO: insular and frontal opercular, MT: medial temporal DSV: dorsal stream visual, Pre: premotor, TPOJ: temporo-parieto-occipital junction, PLMC: paracentral lobular and mid cingulate IF: inferior frontal, AA: auditory association, LT: lateral temporal, EV: early visual).

### D. Hyperbolic embedding of MEG brain networks

#### 1) Hyperbolic Geometry

Hyperbolic spaces are non-Euclidean spaces characterized by negative curvature (*K* < 0) as opposed to zero curvature in Euclidean spaces and positive curvature in spherical spaces. They are represented through various models, prominently the Poincaré disk model and the Lorentz (also known as hyperboloid) model, each offering distinct insights into their geometry and properties (Fig. 2). Recently, hyperbolic spaces have gained traction in representing tree-like graphs due to their capacity to represent hierarchical data effectively [27]. For instance, in a Poincaré model of hyperbolic space, a binary tree can be embedded with minimal distortion (see Fig. 2b).

**Fig. 2.**
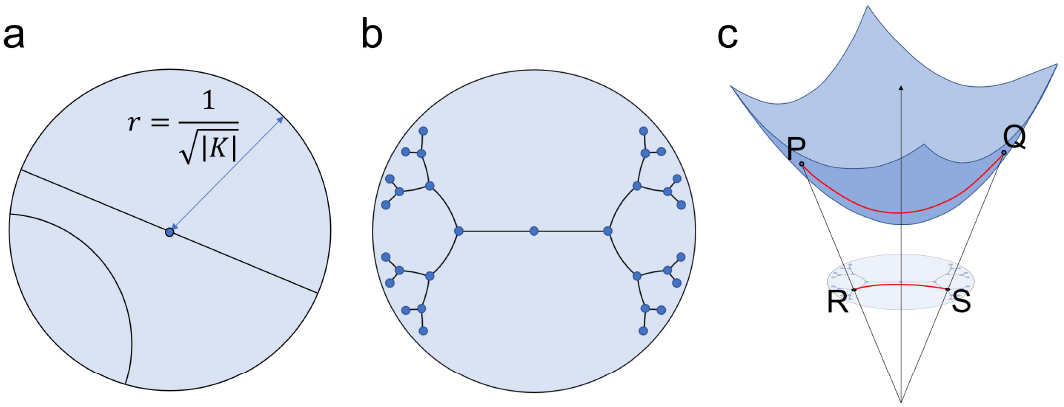
Illustration of Poincaré and Lorentz hyperbolic models. a) Poincaré space, represented as an ***n***-dimensional open ball of fixed radius 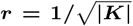 Geodesic lines are arcs bent towards the origin due to the negative curvature. b) An example of a binary tree embedded in the Poincaré space, c) Lorentz space, isomorphic to Poincaré space, depicted here with a projection onto a 2-dimensional Poincaré Disk. The projection mapping function acts as a generalization of stereographic projection to hyperbolic space.

Due to numerical instabilities inherent in the Poincaré model distance metric, recent studies have advocated the use of the Lorentz model [28], [27], [11]. In alignment with these works, we adopt the Lorentz model in our study. Geodesic distance in the Lorentz space can be computed by:

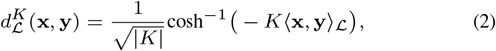

where *K* < 0 is the hyperbolic space curvature, and ⟨**x, y**⟩_ℒ_is the Lorentzian inner product between points **x** and **y**.

Many essential operations in hyperbolic neural networks pose challenges, either being computationally intensive or lacking clear definitions within hyperbolic space. Consequently, these operations are typically handled through a hybrid approach. This involves translating features between the hyperbolic space 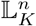 and a tangent space 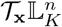 (a Euclidean subspace) around a hyperbolic point 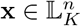 and executing the operations within this tangent space. The transition between spaces is facilitated by logarithmic and exponential maps. Specifically, the logarithmic map, 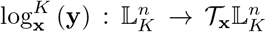 maps a point 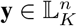 to the tangent space 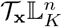

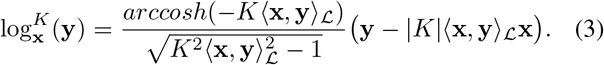

Conversely, the exponential map 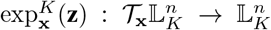 plays an opposite role, lifting a point 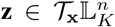 from the tangent space to the hyperbolic space 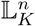

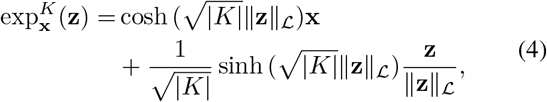

where 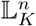 is the n-dimensional Lorentzian hyperbolic space (hyperboloid model) with negative curvature 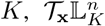 is the tangent space centered at 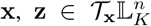, and 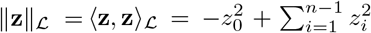. Studies typically set 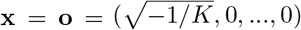 to establish exponential and logarithmic maps between hyperbolic space and the tangent space at the origin.

#### 2) Fully Hyperbolic Neural Network

Previous hyperbolic neural network models primarily formalized operations within intermediate tangent spaces, making them not fully hyperbolic. The logarithmic and exponential maps required to transition to the tangent space involve a series of hyperbolic and inverse hyperbolic functions, whose compositions are complex and often extend to infinity, which significantly undermine the stability of the models. However, recent advancements have led to the development of fully hyperbolic neural networks (FHNN) [15]. This innovation was achieved by relaxing certain restrictions in Lorentz transformations that complicate computations and optimizations, and by designing network layers with feature transformations that inherently preserve data within the Lorentz space. This Lorentz model variant makes it possible to directly formalize operations in hyperbolic space, obviating the need for intermediate tangent spaces via exponential and logarithmic maps. Additionally, the Lorentz model, although harder to visualize, is more numerically stable than the Poincaré model [28].

Further, to formalize operations such as attention and aggregation directly in hyperbolic space, the FHNN model leverages a proof by Law et al. [28], which shows that, with squared Lorentzian distance defined as 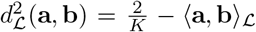, the centroid induced by this distance has a closed-form given by:

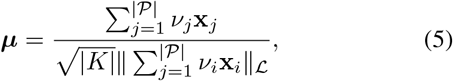

where 𝒫 is the point set 𝒫 = **x**_1_, …, **x**_|𝒫|_, with **x**_*i*_ being the *i*-th node’s hyperbolic embedding representation and *ν*_*I*_ being its weight. This calculation is much faster than current algorithmic implementations like the Fréchet mean in standard hyperbolic spaces [29] [30]. The FHNN centroid is also better than the Einstein midpoint since the latter requires mapping back and forth between the Klein and Poincaré models in order to compute the centroid, potentially leading to information loss [15], [31].

Our adapted FHNN architecture is illustrated in Fig. 3 and can be summarized into three parts: feature transformation, neighborhood aggregation, and non-linear activation. The feature transformation corresponds to a Lorentz linear layer, while the closed-form centroid of neighboring node features defined before is used for neighborhood aggregation. Non-linearity is integrated into the Lorentz linear layer without the need for an explicit non-linear activation function. Below we discuss the model’s individual components.

**Fig. 3.**
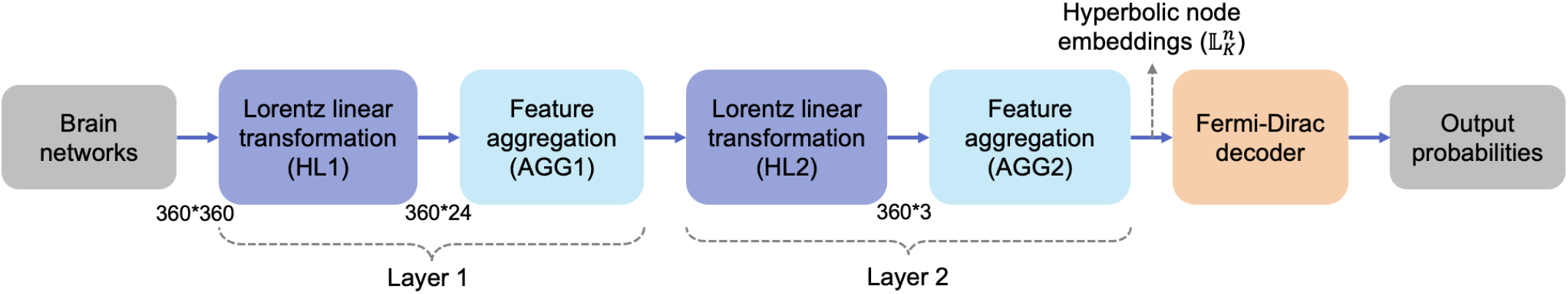
Main architecture of the FHNN model for the link prediction task in MEG brain networks. Each input high-dimensional MEG brain network of size **360** × **360** is first projected to a low-dimensional hyperbolic Lorentzian manifold (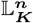 where ***K* < 0** is the curvature, ***n*** is the dimensionality of manifold) with learned “continuous hierarchies” through two consecutive projection layers with output feature sizes **360 × 24** and **360 × 3**, respectively. Each projection layer contains two main operations: hyperbolic (or Lorentz) linear transformation (HL) and feature aggregation (AGG). The Fermi-Dirac decoder is then used to decode the edge probabilities for link prediction from the learned node hyperbolic embeddings.

#### Model Input Features

In graph neural networks, node features capture additional information associated with each node in a graph. In absence of other relevant information, we used a common approach: one-hot identity vectors.

#### Fully Hyperbolic Linear Layer – Feature Transform

The hyperbolic linear layer was devised to automatically preserve data in the Lorentz space, and includes dropout, bias and normalization:

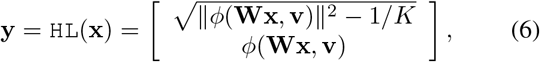

where 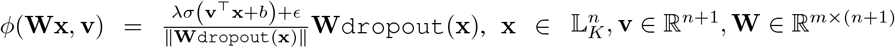 *σ*is the sigmoid function, b’ is a bias term, and *λ >* 0 controls the scaling range.

#### Fully Hyperbolic Aggregation Layer

Relying on the centroid induced by the squared Lorentzian distance [28] (Eq. 5), the aggregation layer in the FHNN uses the following equation:

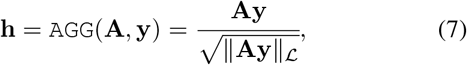

where **A** denotes the adjacency matrix of the MEG brain network, and **y** is the layer input.

#### Link Prediction

Our FHNN model was trained using a link prediction task, which is a common evaluation method in network science [27]. In this task, we aim to predict the presence or absence of edges within a network. Specifically, the total edges of the MEG brain networks were divided into three subsets: a training set (85%), a validation set (10%), and a test set (5%). Non-edges were generated to match the number of edges in the validation and test sets. To perform link prediction, we masked a set of edges from the graph and trained the model to predict whether each edge should exist. The model’s performance was evaluated based on its ability to correctly predict these connections. The Fermi-Dirac decoder translated the proximity of node embeddings in hyperbolic space into probabilities for edge existence [27]. The probability score of a link (or edge) between two nodes *i* and *j* at the output layer *L* is defined as:

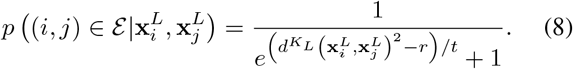

That is, an edge between two nodes is predicted to exist according to a sigmoidal transformation of the hyperbolic distance between the node embeddings. The hyperparameters *r* and *t* control the inflection point and steepness of the sigmoid function, and were set as *r* = 2 and *t* = 1. Finally, the link prediction loss function employed in our model is a margin loss:

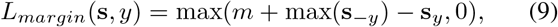

where *m* is the margin, **s** is the output vector of predicted probabilities or scores for each category (in this case, there are two categories: edge exists vs. does not), *y* is the label, **s**_*y*_ is the component of the correct label, and **s**_−*y*_ is the vector composed of all other components corresponding to categories other than the label.

### E. Parameter settings and model training

We divided the entire dataset of 587 MEG connectivity matrices into three sets: 70% for training, 20% for validation, and 10% for testing. To obtain the hyperbolic node embeddings, we trained the FHNN model on a link prediction task [15], which enabled the model to learn both the local and global geometric information of the MEG brain networks. The model was trained with a margin loss function (Eq. 9).

Graph neural networks, including FHNNs, often benefit from shallower architectures to prevent over-smoothing, where node representations become indistinguishable after aggregating information from too many layers. In our case, the small size of the 360 × 360 brain graphs allowed us to use two layers without over-smoothing (Fig. 3). The input features, initially sized 360 × 360, were reduced to 360 × 24 in the first layer and then to 360 × 3 in the second layer. The choice of a smaller embedding size (24) minimized overparameterization while maintaining model accuracy in link prediction. An output embedding size of 3 was selected based on previous studies, which indicated that larger embedding sizes lead to diminishing returns in link prediction performance while increasing the costs of interpretability and overparameterization [29]. We normalized the features and row-normalized the adjacency matrix.

Graph neural networks (GNNs), including FHNNs, often benefit from shallower architectures to prevent oversmoothing, where node representations become indistinguishable after aggregating information from too many layers. In our case, the small size of the 360×360 brain graphs allowed us to use only two layers without over-smoothing: a 360×24 layer and a 24×3 layer. The choice of a smaller embedding size (24) minimizes overparameterization while maintaining model accuracy in link prediction.

The parameters of the FHNN consisted of the weights and biases for the feature transformation layer, as defined in Equation 6. In the first hyperbolic layer, the weight matrix had dimensions 360 × 24, with a corresponding bias vector of size 24 × 1. In the second layer, the weight matrix was 24 × 3, and the bias vector was 3 × 1. All weight parameters were randomly initialized using Xavier initialization to provide an optimal starting point for training.

The hyperparameters of the FHNN include the number of epochs (300 max), early stopping criterion (150 epochs), gradient clipping (0.1), learning rate (0.025), weight decay (0.001), and batch size (64). We used a curvature value of c = 1.0, Fermi-Dirac encoder values of r=2 and t=1, and a margin of 2 for our link prediction margin loss function. The choice of hyperparameters were based on a combination of prior studies [15] and grid search to optimize performance. Training was performed using the Riemannian Adam optimizer.

## III. Results

### A. Alterations of brain network hierarchy across age

Hyperbolic embeddings are powerful tools for representing hierarchical data due to the natural alignment between the geometric properties of hyperbolic space and the structural characteristics of hierarchies. In hyperbolic space, hierarchical relationships become more pronounced: central nodes often represent higher-level categories, while peripheral nodes correspond to subcategories. This is illustrated in Fig. 2b, where nodes near the root of the tree are placed close to the origin, and nodes further from the root are situated farther out. Leveraging this property, we calculated the hyperbolic radius of each brain region as an indicator of its position within the overall brain hierarchy. This metric, computed for each brain region across the 587 participants of the Cam-CAN database, allowed us to characterize subtle hierarchical organizational changes in various brain subnetworks associated with the aging process. In our study, we used the van Essen HCP multimodal atlas, which comprises 360 brain regions that were subsequently grouped into 22 subnetworks, organized according to their structural properties, functional task-related profiles, and functional connectivity patterns [24].

We estimated the average hyperbolic radius for brain regions within each of 22 brain subnetworks and across participants of the same decade (Table I), as depicted in Fig. 4. The findings suggest a consistent increase in the hyperbolic radius across most of the 22 brain subnetworks over the decades, indicative of a diminishing hierarchical significance over time. Notably, certain subnetworks exhibit more rapid changes over time than others.

**Fig. 4.**
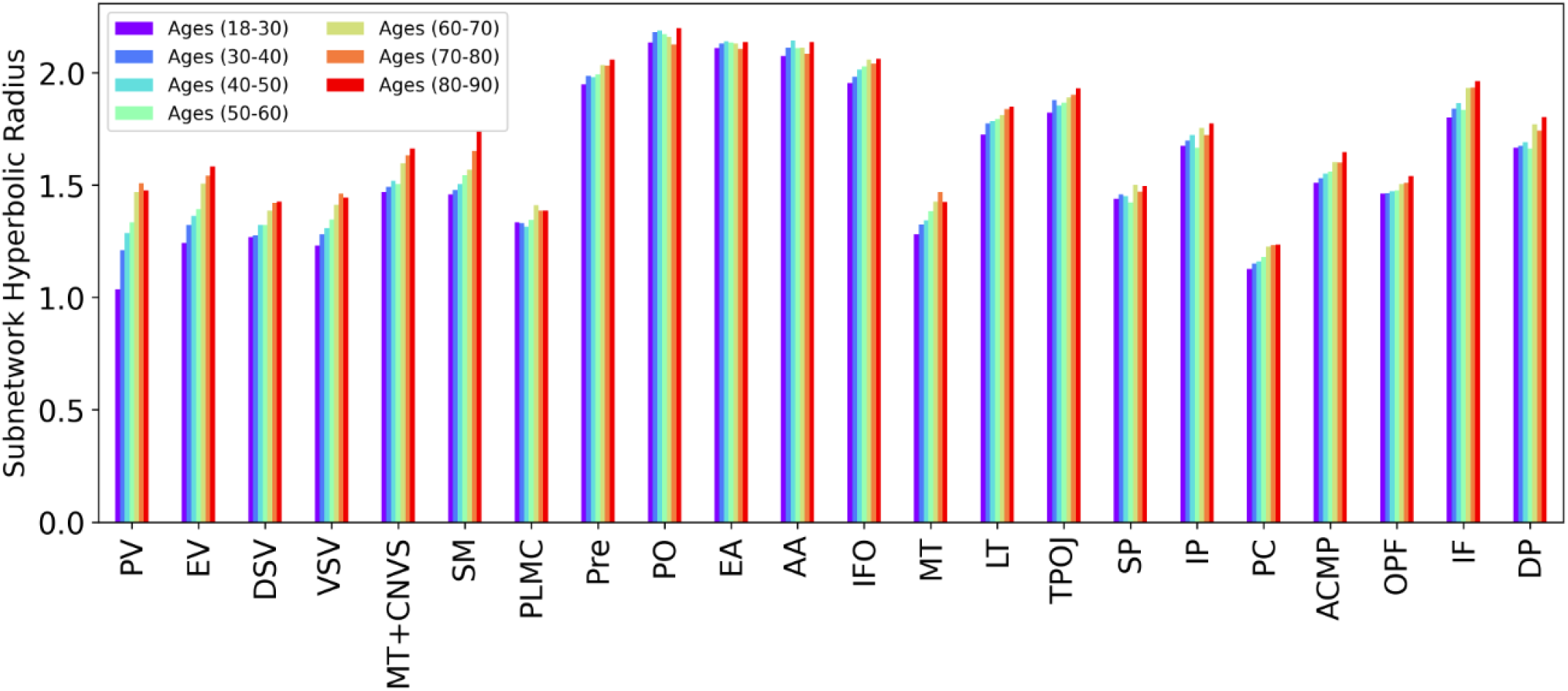
Hyperbolic radius changes for each of the 22 brain subnetworks across 7 age decades. The full subnetwork names are provided in Fig. 1.

To investigate further, we fitted a linear regression model to the 7-decade hyperbolic radius data points for each brain subnetwork, obtaining a slope value that indicates the alteration of the subnetwork’s hyperbolic radius across age (Fig. 5). Our findings reveal statistically significant increases in hyperbolic radius within the majority of the brain subnetworks. Notably, primary cortical areas, including the primary visual, early visual, somatosensory, and motor subnetworks, exhibit the most prominent changes. Conversely, several other subnetworks demonstrate more gradual alterations, with brain subnetworks such as early auditory and posterior opercular exhibiting near-zero slopes across age decades. Additionally, we observed that brain subnetworks with a smaller overall hyperbolic radius tend to have higher slopes, indicating a faster reduction in hierarchy within regions that initially have a high hierarchical organization.

**Fig. 5.**
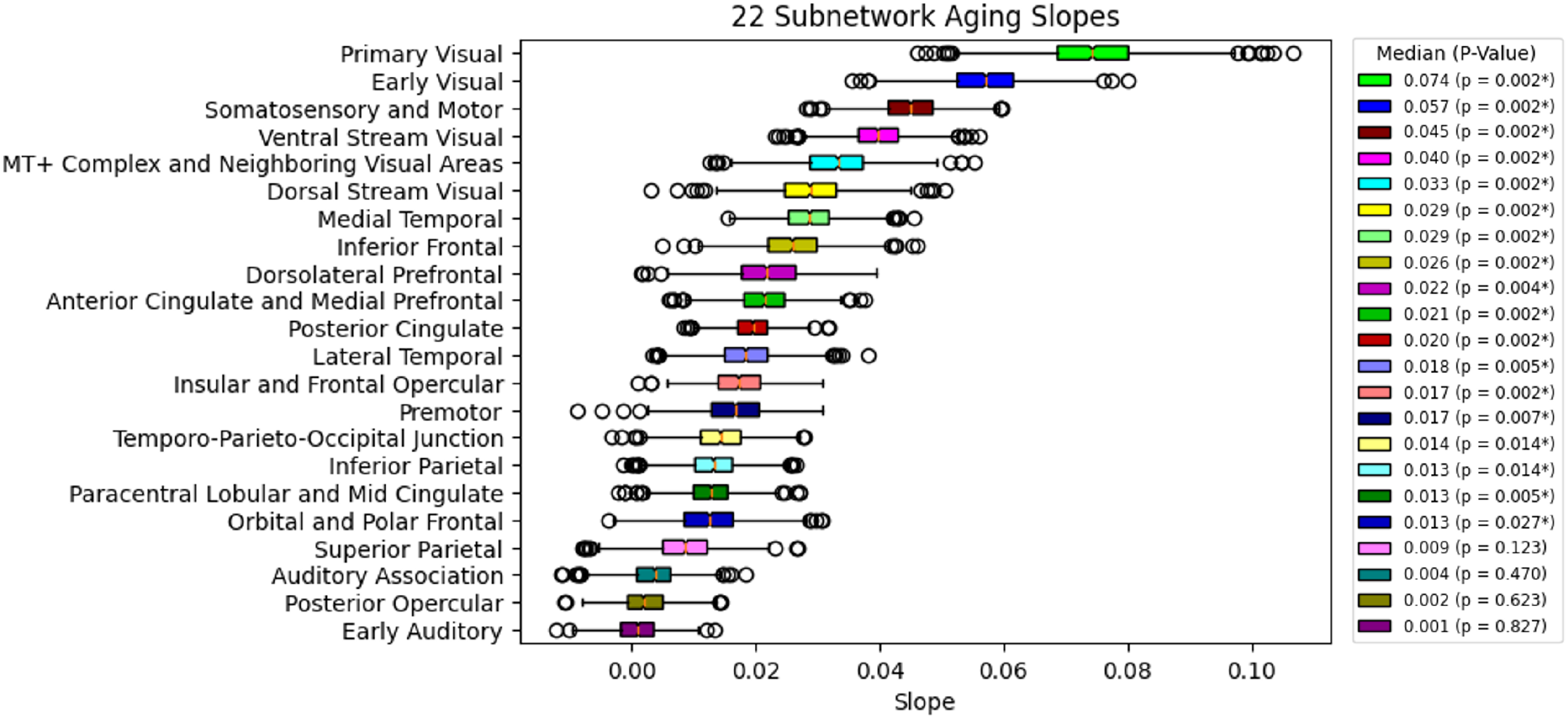
Slope of hyperbolic radius change for each of the 22 brain subnetworks. Each slope was computed by fitting a linear regression model to the 7-decade hyperbolic radius data of the corresponding subnetwork. Box plots were constructed using the distribution of 1000 bootstrap samples from the 587 Cam-CAN participants. P-values were calculated using 1000 permutation samples by randomly shuffling the subject’s age labels and recomputing the slopes for each permutation sample. These p-values were then adjusted for multiple comparisons using false discovery error rate. Statistically significant p-values at a 5% level are indicated with a star (*).

The slopes of the age-related hyperbolic radius changes are visualized on a smoothed cortical manifold for the 360 brain regions comprising the HCP-MMP1 atlas [24] (Fig. 6). Additionally, we illustrate the hyperbolic embeddings of two brain subnetworks with contrasting aging trajectories in Fig. 7. The embeddings of brain regions within the early visual subnetwork, which exhibits a high slope, are shown for both the 1st decade (18-30 years) and the 7th decade (80-90 years). In the 7th decade, these embeddings have shifted towards the periphery, reflecting a pronounced increase in the overall hyperbolic radius and thus a decrease in brain network hierarchy. For contrast, we also present the hyperbolic embeddings of areas within the early auditory subnetwork, a region with a nearly flat slope and stable hyperbolic radius from the 1st to the 7th decade. Note that the hyperbolic embeddings are isomorphically projected from the Lorentz space, the native space of the FHNN model, to the Poincaré disk for easier visualization.

**Fig. 6.**
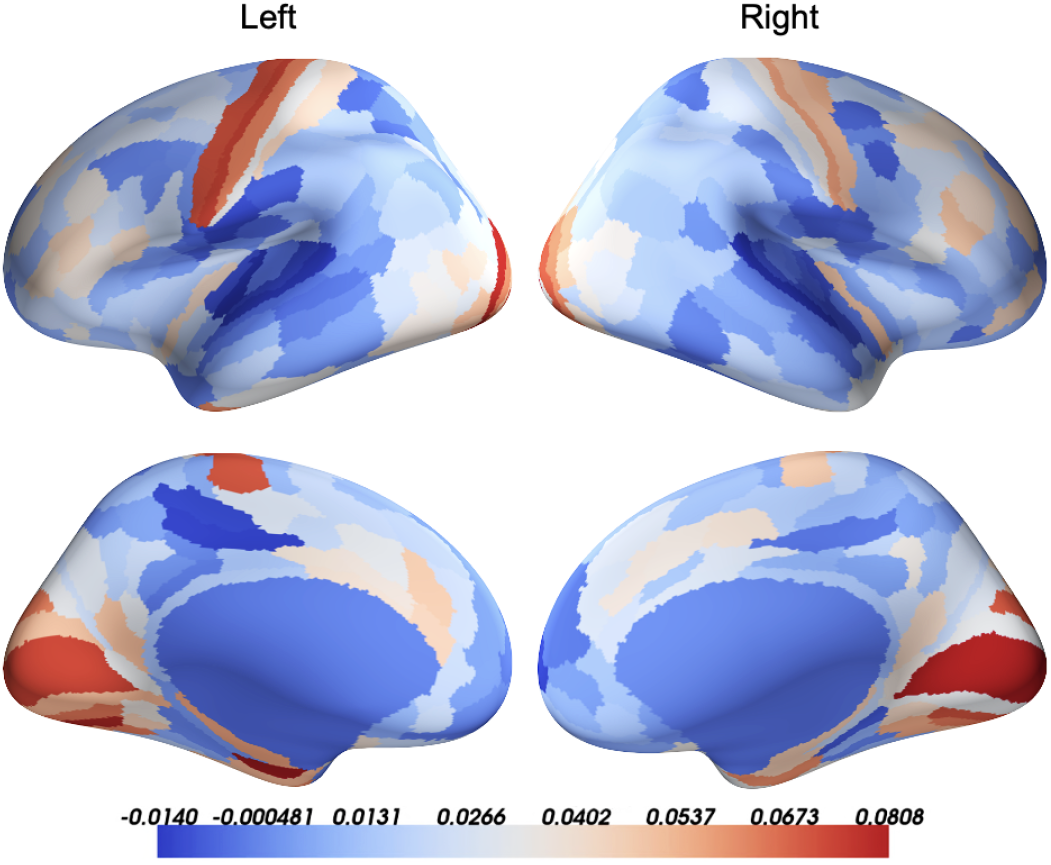
Visualization of age-related slopes of hyperbolic radius change for the 360 brain regions defined in the HCP-MMP1 atlas [24].

**Fig. 7.**
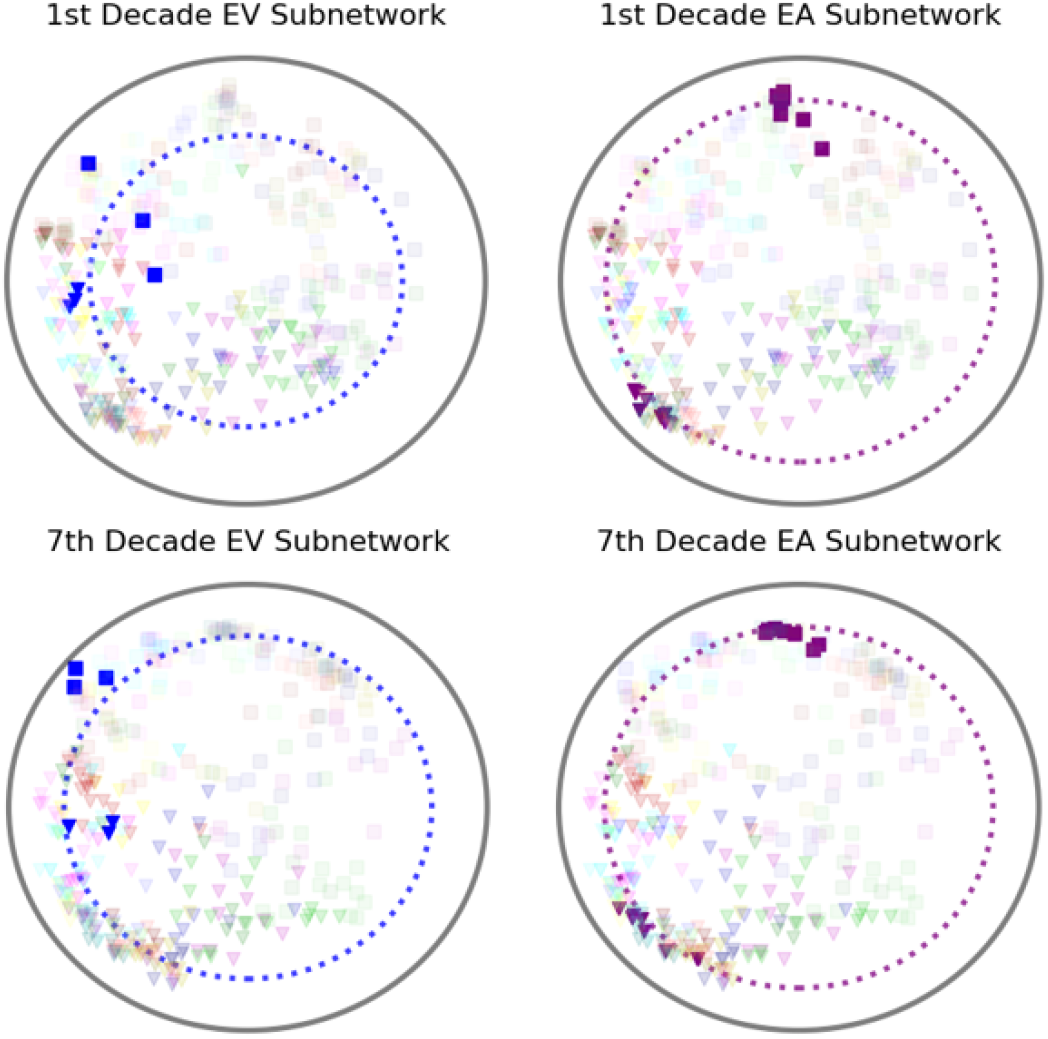
Hyperbolic embeddings of the 360 brain regions for participants aged in the 1st decade (18-30 years) and 7th decade (80-90 years). Two subnetworks, the *early visual* and *early auditory*, are highlighted, while the rest are faded. Early visual areas show significant increases in hyperbolic radius, whereas early auditory areas remain stable over time. Individual points represent different brain regions, with shapes indicating the hemisphere (“□” for right hemisphere, “**△** “ for left hemisphere) and colors indicating subnetwork membership, consistent with Fig. 1.

### B. Age prediction with hyperbolic radius

Deep learning is highly effective at automatically discovering informative features from data for downstream tasks. We hypothesized that hyperbolic radii, which reflect the automatically detected hierarchy of each brain region from the FHNN model, would serve as more powerful features than traditional hand-crafted graph-theoretic features that also reflect node importance, such as closeness centrality and betweenness centrality.

To evaluate, we assessed the effectiveness of using the hyperbolic radius of the 360 brain regions versus standard graph-theoretic measures for the downstream task of age prediction. For comparison, we computed eight commonly used standard graph-theoretic measures—shortest path length, efficiency, clustering coefficient, degree centrality, Katz centrality, closeness centrality, betweenness centrality, and average neighbor degree—for the MEG brain networks of the 587 subjects. To ensure a robust and comprehensive comparison, we tested three regression methods (AdaBoost, Ridge Regression, and Random Forest) with features consisting of the hyperbolic radii or the standard graph measures. We also compared performance when concatenating both features. Model fitting was performed using the same 70/30% train-test split as in the FHNN training procedure to ensure generalizability.

The corresponding age prediction performance, measured by Mean Absolute Error (MAE), is shown in Table II. We observe that the hyperbolic radius outperformed all standard graph-theoretic measures, achieving lower MAE across all the three different regression models. Further, by concatenating hyperbolic radius as an additional regression feature to the standard graph measures, all three regression models improved performance relative to using standard graph measures alone. Our findings validate that hyperbolic node embeddings are rich low-dimensional representations of MEG brain networks that encode more powerful age-related information in brain networks than standard graph measures.

The implementation of our adapted FHNN model for MEG brain network embeddings can be found in [link removed for double blind review].

**TABLE 2.** Mean Absolute Error (%) Results for Age Prediction with Different Regression Models and Features (the MAE differences between applying individual measures and combined each feature with hyperbolic radius feature are shown in parentheses.)

| Graph Measure Feature | Individual Measures |  |  | Measures Combined with Hyperbolic Radius |  |  |
| --- | --- | --- | --- | --- | --- | --- |
|  | Ada-Boost | Ridge | Random Forest | Ada-Boost | Ridge | Random Forest |
| Shortest Path Length | 15.24 | 14.83 | 15.36 | 13.16 (-2.08) | 13.19 (-1.64) | 13.29 (-2.07) |
| Efficiency | 14.07 | 13.81 | 14.03 | 13.05 (-1.02) | 13.26 (-0.55) | 13.17 (-0.86) |
| Clustering | 13.91 | 14.92 | 14.23 | 12.74 (-1.17) | 13.08 (-1.84) | 12.99 (-1.24) |
| Degree Centrality | 13.85 | 13.62 | 13.92 | **12.73** (-1.12) | **13.12** (-0.50) | **12.78** (-1.14) |
| Katz Centrality | 15.29 | 15.88 | 15.24 | 13.57 (-1.72) | 13.77 (-2.11) | 13.31 (-1.93) |
| Closeness Centrality | 14.10 | 14.05 | 14.37 | 12.9 (-1.20) | 13.30 (-0.75) | 12.99 (-1.38) |
| Betweenness Centrality | 14.08 | 15.58 | 14.10 | 12.95 (-1.13) | 13.4 (-2.18) | 12.79 (-1.31) |
| Average Neighbor Degree | 14.48 | 14.19 | 14.50 | 12.76 (-1.72) | 13.42 (-0.77) | 12.96 (-1.54) |
| Hyperbolic Radius | **13.01** | **13.16** | **13.00** | — | — | — |

## IV. Discussion

We developed a novel hyperbolic MEG brain network embedding framework based on the FHNN architecture [15], and proposed using the hyperbolic radii of the node embeddings as features to study aging trajectories. Our findings revealed that hyperbolic embeddings are powerful signatures of the aging brain, surpassing traditional graph theoretic measures in capturing age-related information from MEG brain networks.

Our findings revealed that a significant number of subnetworks underwent a reduction in hierarchy during aging, with some exhibiting gradual changes while others experienced rapid transformations. Similar trends have been previously observed using graph-theoretical measures on resting state Fmri [32] and MEG data [33]. Despite its potential relevance as a diagnostic marker for neurodegenerative disorders, the general trend of brain-network reorganization during healthy aging and its dependence on specific brain subnetworks remains poorly understood. Our novel metric, directly related to the loss of hierarchical organization as quantified by the hyperbolic radius, may offer greater efficiency and interpretability compared to classical graph-theoretical measures.

While the most pronounced changes in the slope of the hyperbolic radius were concentrated in the visual, somatosensory, and motor areas, we also observed significant positive slopes across a wide range of brain regions. Notably, this included prefrontal areas such as the Dorsolateral Prefrontal Cortex (DP) and the Anterior Cingulate and Medial Prefrontal Cortex (ACMP). This aligns with the hierarchical organization of the prefrontal cortex at the network level and its critical role in modulating cognitive functions that change with age.

Our study focused on characterizing normal brain aging trajectories rather than diagnosing neurodegenerative diseases. Understanding typical aging patterns is crucial for identifying deviations that may indicate early pathological changes. Although our study does not include patient data, it provides a foundational methodology for future research aimed at discovering biomarkers for neurodegenerative disorders.

## V. Conclusion

Our hyperbolic MEG brain network embedding framework effectively transformed complex high-dimensional MEG brain networks into lower-dimensional hyperbolic representations, facilitating brain hierarchy analysis across age. Our findings revealed that a considerable number of brain subnetworks exhibited a reduction in hierarchy during aging.

In contrast to other machine learning models, our methods generate hyperbolic node embeddings that offer straightforward interpretation through visualization in the Poincaré Disk. These embeddings carry tangible meaning, demonstrated by the hyperbolic radius serving as a measure of hierarchical brain organization. Consequently, one can readily distinguish abnormal brain networks from normal ones with greater interpretability compared to models lacking a direct association between internal representations and physiological significance. Overall, our study presented the first evaluation of hyperbolic embeddings in MEG brain networks across age, introduced a novel neural embedding measure of brain hierarchy, and used this measure to highlight aging trajectories in the large cohort of the Cam-CAN dataset. Our approach has the potential to deepen our understanding of the aging brain, help in delineating normative brain aging trajectories, and pinpoint deviations indicative of neurodegenerative disorders, such as Alzheimer’s disease.

